# Managing Gaze Competition when Acting On and Monitoring the Environment in Parallel

**DOI:** 10.1101/2024.10.28.620734

**Authors:** Jolande Fooken, Roland S. Johansson, J. Randall Flanagan

## Abstract

Research on visually guided object manipulation has shown that participants fixate goal locations-such as objects to be grasped and locations where they are placed-prior to hand arrival, with gaze serving two primary functions: *directing* the hand (or object in hand) to the vicinity of the goal using peripheral vision and gaze related signals, and *guiding* the hand using central vision as it approaches the goal. However, in real world scenarios, manipulation tasks are often performed while concurrent monitoring of the environment, resulting in competition for gaze. Here we examined gaze-hand coordination under such conditions. Participants performed a manipulation task, that involved grasping balls and placing them at target locations, while concurrently monitoring a display to detect probabilistically occurring visual events, which required central vision. Participants managed gaze competition in two main ways. First, fixations allocated to the action task were brief and prioritized *directing* the hand towards the goal (object or target location); participants then relied on tactile feedback to complete the action (grasping or placing the object). When tactile feedback was reduced-by using a tool instead of the fingertips to perform the task-gaze additionally served the *guiding* function. Second, participant reduced gaze competition by exploiting temporal regularities of events in the monitoring task. Specifically, they adjusted both gaze allocation and hand movement timing to reduce the likelihood that action task fixations would coincide with visual events. These findings demonstrate how individuals flexibly integrate sensorimotor control with analysis of environmental statistics to manage competing visual demands.

**Significance Statement:** In everyday behaviour, we often perform manual tasks while simultaneously monitoring the environment, creating competition for gaze. How the brain resolves this competition remains poorly understood. Using a novel paradigm combining an object manipulation task with visual event monitoring, we show that participants integrate knowledge of sensorimotor demands and temporal regularities in the monitoring task to manage gaze. Specifically, we found that participants preferentially allocated gaze to the action task when it is most critical for sensorimotor control and when the likelihood of a visual event was low. Additionally, participants adjusted their hand movement timing based on event statistics to reduce gaze competition. These findings reveal how the brain dynamically allocates gaze resources across competing sensorimotor and visual task demands.

## Introduction

In everyday life, we often perform manual tasks while also monitoring the environment for events of interest, leading to competition for gaze between action and observation (Land and Furneaux, 1997; Kowler, 2011; Hayhoe, 2017; Fooken et al., 2023; Keshava et al., 2024). For example, while cooking, one may use gaze both to slice vegetables and to detect the moment when water begins to boil or a pan starts to smoke. In principle, we can consider three strategies by which individuals might manage this competition. First, individuals may selectively allocate gaze to the action task only when gaze is most critical for sensorimotor control. Second, they may exploit temporal regularities of events in the visual environment, preferentially directing gaze to the action task when the likelihood of an event is low. Third, they may further leverage these regularities by modifying the timing of their manual actions so that gaze-critical phases of the task coincide with periods of reduced event probability.

To investigate how people manage competing demands on gaze, we designed a novel paradigm (Fig. 1A) in which participants performed a continuous object manipulation task while monitoring a display to detect visual events (letter changes). The manipulation task involved grasping a small ball, moving it to a slot, and inserting it, and thus captured the key components of many manual actions: grasping, transporting, and placing objects (Johansson et al., 2001; Flanagan et al., 2006). Visual events in the monitoring task occurred probabilistically, with the likelihood of events varying predictably over time.

**Figure 1.**
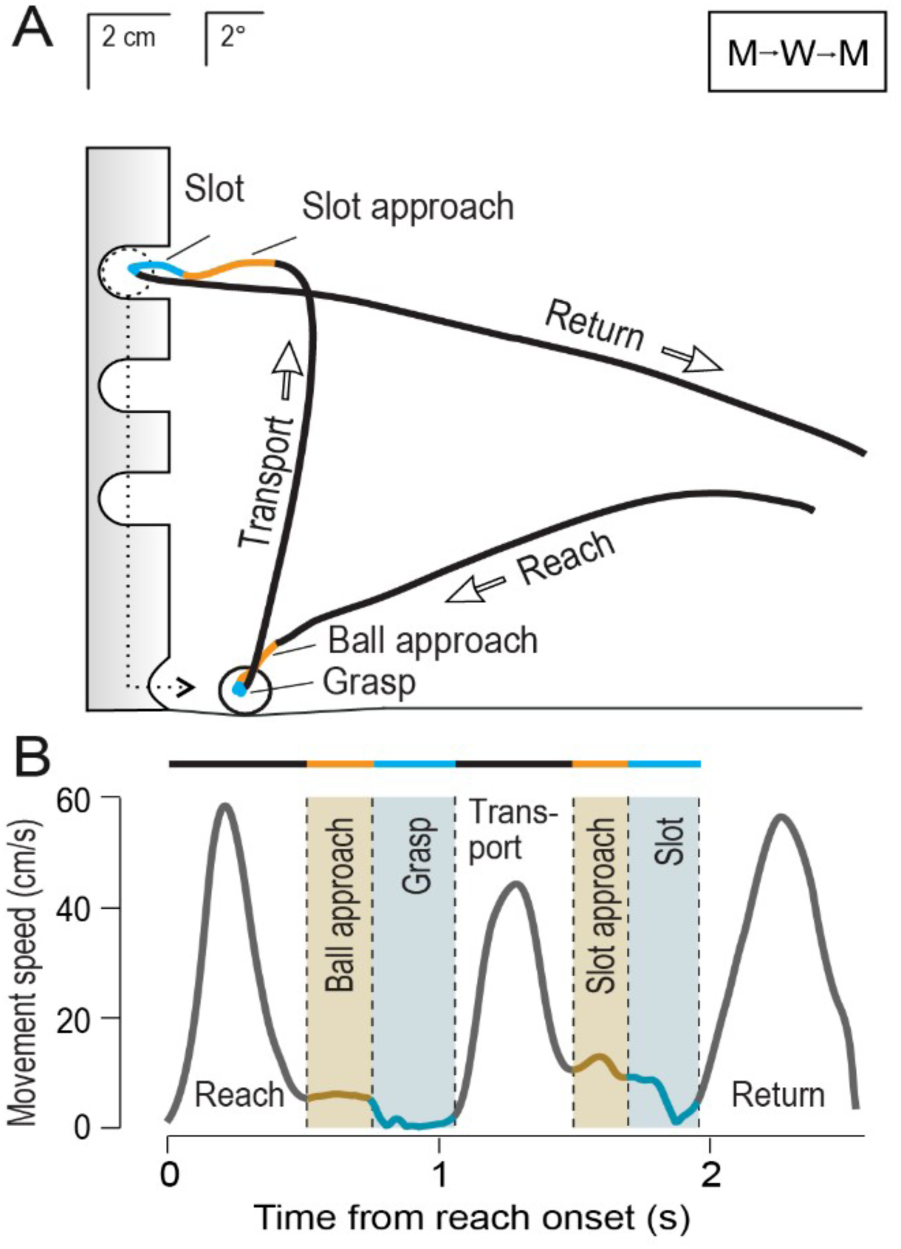
Apparatus and action phases in the ball-drop task. (A) Participant’s view of the task setup, showing the ball-drop platform and text display (top right), and the hand movement path in a single trial. Movements were executed in a frontal plane positioned ∼40 cm in front of the participant’s eyes. The vertical tube contained three slots. (B) Representative end-effector velocity profile from a single trial, illustrating the segmentation of the trial into 7 action phases based on kinematic and contact events (see Materials and Methods for details). These phases include: (1) Reach-movement initiation from the home position; (2) Ball approach-deceleration toward the ball; (3) Grasp-ball contact to lift-off; (4) Transport-movement toward the slot; (5) Slot approach-final deceleration before insertion; (6) Slot-entry into the slot and ball release; and (7) Return-movement back to the starting position.

When manual tasks are performed without competing visual demands, gaze typically shifts between action targets and can serve three primary functions: *directing* the hand movement toward the vicinity of the current target using peripheral vision and gaze related signals (Goodale et al., 1986; Saunders and Knill, 2003, 2004), *guiding* the hand with central vision as it nears the target (Ballard et al., 1992; Johansson et al., 2001; Land, 2006), and *checking* goal completion (e.g., confirming an object has been successfully grasped) also using central vision (Safstrom et al., 2014). We hypothesized that in the face of gaze competition, participants would prioritize the *directing* function, relying primarily on tactile feedback from the fingertips to complete the action goal (e.g., grasp the object). In a condition in which participants used a tool (tweezers) to perform the manipulation task, we further predicted that both the *directing* and *guiding* functions would be prioritized, due to reduced tactile feedback and increased precision demands.

Previous research has shown that people can learn and exploit temporal regularities of events to optimize gaze allocation when monitoring multiple locations (Hoppe and Rothkopf, 2016). We predicted that participants would similarly leverage the temporal structure of events to reduce gaze competition while concurrently manipulating objects and monitoring the environment.

Specifically, we hypothesized that participants would be more likely to fixate action targets (i.e., the ball and slot) during periods of low event probability. Additionally, we hypothesized that participants would adjust the timing of their hand movements to lower the likelihood of an event occurring at moments when gaze is most critical for the action.

Confirmation of these hypotheses would provide evidence that people can flexibly integrate sensorimotor control with temporal prediction based on environmental statistics to adaptively allocate gaze when simultaneously manipulating and monitoring the world.

## Methods

### Participants

Eleven right-handed participants (8 male; age 22-33 years) with normal or corrected-to-normal vision took part in the study. All were naive to the study purpose. Written informed consent was obtained prior to participation. The study was approved by the ethics committee at Umea University.

### Apparatus and general procedure

Participants were seated at a table with the ball-drop apparatus positioned in front of them (Fig. 1A). It consisted of a 15 cm high vertically oriented Perspex tube (inner diameter = 14 mm; wall thickness = 3 mm) mounted on a wooden platform. The tube was positioned approximately 2.5 cm to the left of the participant’s mid-sagittal plane, with its top aligned at eye level. Three vertical slots were cut into the front of the tube, with their centers located at 5, 8, and 11 cm above the platform surface.

Participants continuously performed the ball-drop task, which required them to reach for and grasp a small ball (12 mm diameter polished brass sphere) at a designated start position on the platform, transport it the target slot, insert it into the tube, and release it. After it was released, the ball dropped through the tube and rolled back to its starting position on the platform via a slightly sloped surface. One second after the ball returned to the start position (or was already present at the start location in the first trial), a pre-recorded auditory instruction (“bottom,” “middle,” or “top”) indicated the slot to be used (i.e., the target slot) in the upcoming trial.

All movements were performed in a vertical plane aligned with the participant’s coronal plane and located 40 cm from the eyes. The start position for the ball was located 3 cm to the right of the tube’s vertical midline. After releasing the ball, the participant returned their hand to a support plate-the hand start position in each trial-positioned 20 cm to the right of the platform. In separate trial blocks, the ball was grasped either directly with the fingertips or with a handheld tool (plastic tweezers, 14 cm long, with 4 mm diameter tips coated with thin plastic tubing to enhance friction). Note that we included trial blocks with this tool to increase the visuomotor demands of the action task. This increase can be attributed to three factors. First, the contact surfaces of the tool tips are small compared to the fingertips and thus require greater spatial precision, particularly during ball grasping. Second, unlike the more malleable fingertips, the rigid tool tips do not conform to the ball’s surface, reducing grasp stability. Indeed, in robotic manipulation, soft contact surfaces are used to enhance stability and reduce spatial precision demands (Bicchi, 2000; Billard and Kragic, 2019). Third, the tool provides limited tactile feedback about the contact state, and impaired tactile sensibility is known to increase reliance on vision during object manipulation (Brink and Mackel, 1987; Jerosch-Herold, 1993; Jenmalm and Johansson, 1997; Chemnitz et al., 2013).

In the single task conditions, participants performed the ball-drop task only. In the dual task conditions, participants concurrently performed a visual monitoring task that involved detecting brief letter changes (LCs) on an LED text display, which required central vision. The text display was positioned in the upper right quadrant of the scene (Fig. 1 C and D), 7 cm behind the frontal plane in which the tube was located. The visual angle between the center of the letter area and the center of the top slot and ball were 24° and 28°, respectively.

The letter height and width corresponded to 0.5° x 0.7° visual angle. At randomly distributed intervals (1.5-6.5 s, uniform distribution), the letter M was changed to W for 300 ms before reverting to M. The participants were instructed to press a button held in their left hand as soon as they detected a LC. If they failed to respond within 1 second after a change, the LC was considered missed. Missed detections were signaled by a brief beep and flashing hash marks on the display for 600 ms. To ensure that the participants relied on central vision to detect LCs rather than peripheral visual cues, the letter M randomly shifted its horizontal position by 0.6° visual angle at intervals between 1 and 3 s (uniform distribution). Pilot tests confirmed that participants had to foveate the display to detect LCs.

To ensure that participants remained engaged with both the object manipulation task and the monitoring task, a performance-based incentive system was implemented. Participants earned 1 Swedish krona (SEK) for each successful ball-drop and incurred a penalty of 3 SEK for each missed letter change (LC).

Each participant completed four conditions in a fixed order: single-task with fingertips, single-task with the tool, dual-task with fingertips, and dual-task with the tool. Each condition included 30 trials (10 per slot) with the order of target slot varying unpredictably across trials. Note that each participant completed three versions of the experiment, that differed in how the correct slot was specified. In the version reported here, the slot was cued auditorily; in the visual cue version, LEDs on the tube indicated the slot; and in the ‘memory’ version, participants were instructed to follow a predefined slot sequence from memory. The order in which these three versions were completed was counterbalanced across participants. As no clear differences in manual or gaze behaviour were observed across versions, we decided to focus our analysis on the auditory-cue version.

### Data collection and analysis

Gaze position was recorded at l20 Hz (RK-726PCI pupil/corneal tracking system, ISCAN Inc., Burlington, MA). The head was stabilized with chin and forehead support. The standard deviations of gaze position measurement errors were 0.50° (horizontal) and 0.52° (vertical) of visual angle, corresponding to 0.35 cm and 0.36 cm in the vertical plane in which the ball was moved during the task.

The positions of the right index finger and the tool tips were tracked at 60 Hz using 6 DOF position-angle sensors (RXl-D miniature receiver; FASTRAK, Polhemus, Colchester, VT). The fingertip sensor, attached to the nail, recorded the preferred contact point with the ball. The mapping between the position and orientation of the sensor and the preferred contact points was established during calibration trials in which the participants grasped the stationary ball at a known position. The position of the tool sensor, attached to the proximal end of the tool, was offset to the midpoint between the tool tips.

A six-axis force-torque transducer (Nano F/T transducer, ATI Industrial Automation, Apex, NC; sampling rate 400 Hz), mounted under the platform plate, detected when the ball contacted the platform after being dropped, was first contacted by the fingertips or the tool, and was lifted off the platform. Optical sensors (SG-2BC, Kodenshi, Japan) indicated when the ball was inserted into the tube and when it was at its start position. Gaze and sensor signals were synchronized and sampled at 200 Hz using SC/ZOOM software (Umea University).

To characterize the movement sequence in the ball-drop task we defined seven action phases based on the speed of the fingertip or the tool (Fig. lB). Speed was computed as the vector magnitude of the first time derivatives of filtered horizontal and vertical position signals (2nd order Butterworth low-pass filter, 10 Hz cut-off frequency). The reach phase began when end-effector speed exceeded 2 cm/s as the hand left its starting position, typically showing a bell-shaped velocity profile. The ball approach phase began at the first local minimum or inflection point in the velocity profile following the reach. The grasp phase started upon contact with the ball and ended at lift-off. The transport phase spanned from lift-off to the next local minimum in speed (typically preceding final slot approach), also bell-shaped. The slot approach phase started at the inflection point following transport and was characterized by reduced speed as the end-effector neared the slot. The slot phase started when the end-effector tip crossed into the tube (1 cm right of release point) and ended at ball release, detected by optical sensors. Finally, the return phase began after release and concluded when the hand returned to the start position, again showing a bell-shaped velocity profile.

Gaze fixations were identified based on previously established criteria (Johansson et al., 2001). Fixations were classified into three categories according to their location, each defined as a centroid with a 2.5 cm radius: the ball at its start position, the target slot, and the text display. Note that fixation data were pooled across all slot positions (bottom, middle, and top). To qualify as a fixation, gaze had to remain within a designated zone for at least 100 ms. Notably, practically all fixations occurred within these zones (-95 %). Consistent with previous findings on object manipulation, participants almost never fixated their hand, or the tool in hand, during hand movement.

We used paired t-tests with Holm-Bonferroni correction, Kolmogorov-Smirnov (KS) tests, and multiple linear regression analyses in our statistical analyses, using a < 0.05. All statistical analyses were conducted in R (R Core Team, 2022; www.r-project.org). Trials in which participants dropped the ball after initiating the transport phase were excluded. A total of 64 trials (4.7 %) were removed, with no participant losing more than 7 trials.

## Results

### Gaze-hand coordination in single-task conditions

To establish a baseline for interpreting gaze behaviour under dual-task demands, we first examine gaze behaviour in the single task conditions. The top panels in Figs. 2A and B show representative gaze and end-effector paths during trials performed with the fingertips and the tool, respectively. The bottom panels show the average speed of the tip of the end-effector (top row) and the instantaneous probabilities of gaze fixating the ball (at its start position) and the target slot as a function of time (bottom row). These averages are based on aggregate data from all trials and participants. To preserve phase-specific information while aligning trials temporally, we normalized the duration of each phase in each trial to the median phase duration within each condition.

**Figure 2.**
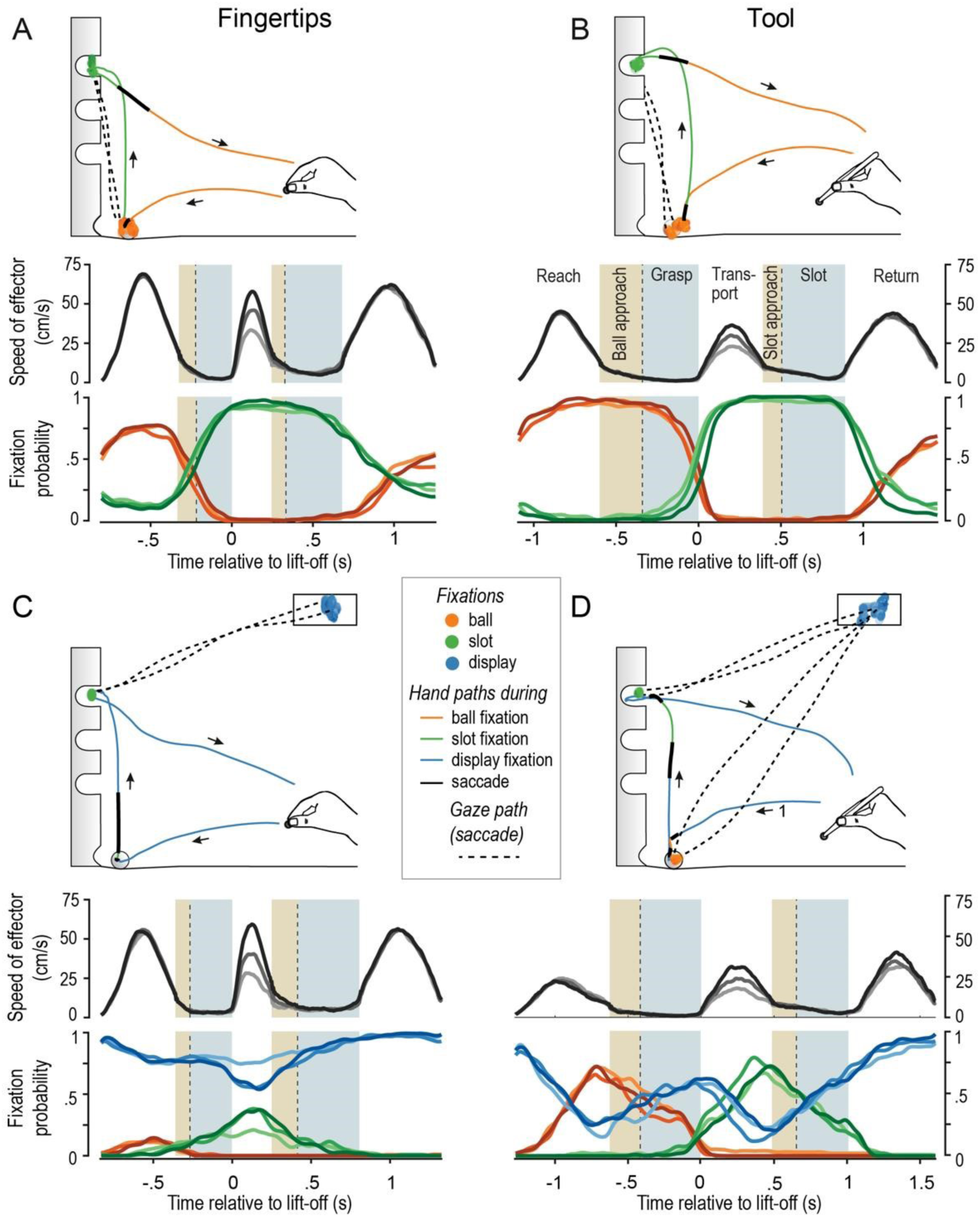
Gaze-hand coordination. (A-D) Top panels show gaze and end-effector paths in single fingertip and tool trials in the single (A, B) and dual task (C, D) conditions. Fixations are color-coded according to landmark, and end-effector paths are color-coded based on the concurrent gaze state. Bottom panels show average end-effector speed and the probabilities of fixating the ball (orange), slot (green), and display (blue) as a function of time relative to lift-off. Separate traces are shown for each slot (top to bottom coded dark to light). Alternating white, brown, and grey-blue regions indicate movement phases labelled in B. Vertical dashed lines mark first ball contact (end of ball approach) and slot entry (end of the slot approach). Data aggregated across all participants, with phase durations in each trial normalized to the median duration of each phase.

In both fingertip and tool trials, participants typically fixated the ball during the reach phase, shifted gaze to the slot around the time ball contact, and maintained the fixation of the slot until around the time the ball was dropped. Gaze then returned to the ball’s start position at the tube’s base. However, in fingertip trials, gaze was occasionally located at the slot during the reach phase. As expected, participants completed the ball-drop trials more rapidly (t_10_ = 4.94, p < 0.001; d = 1.49) when using fingertips (2.04 ± 0.33 s) compared to the tool (2.53 ± 0.31 s). The timing of the gaze shift from the ball to the slot, relative to ball contact, differed between fingertip and tool trials (t_10_ = 13.24; *p* < 0.001; *d* = 3.99). In fingertip trials, the shift occurred just before contact (-0.06 ± 0.05 s; mean ± SE) whereas, in tool trials, the shift occurred well after ball contact (0.29 ± 0.11 s). These findings are consistent with the idea that tool use increases reliance on central vision to establish stable grasp on the ball.

### Gaze-hand coordination when there is competition for gaze

The remainder of the results focus the dual-task conditions. As shown in Figs. 2C and D, participants divided their gaze fixations between the text display and the action-related landmarks. As a result, the probability of fixating the ball or slot during the ball-drop task was reduced compared to the single task conditions, regardless of end-effector. Additionally, the durations of action-related fixations were consistently shorter.

In fingertip trials, participants primarily fixated the display during the reach, ball approach, and grasp phases (88% of trials), indicating that grasping the ball generally did not require central vision. When the ball was fixated, the fixations occurred during the reach phase, with gaze typically shifting away before the ball approach phase, and almost never remaining on the ball after contact. Transporting the ball and inserting it into the slot were also frequently performed with gaze remaining on the display. However, brief fixations of the slot occurred in about half the trials (51%) (see example in Fig. 2C, top panel). The probability of fixating the slot peaked midway through the transport phase and then declined steadily, approaching zero by the end of the slot phase (Fig. 2C, bottom panel). These findings indicate that in fingertip trials, inserting the ball into the slot often did not require central vision.

Consistent with the fixation probabilities described above, we observed two predominant gaze patterns in fingertip trials: ‘display-only’, where gaze remained on the display throughout the trial, and ‘slot’, where gaze shifted from the display to the slot and then back to the display (Fig. 3). To examine the relationship between gaze pattern and manual performance, we compared the action phase durations in display-only and slot trials. Only participants (*N*=9) who demonstrated both fixation patterns were included. The duration of the slot phase was shorter (*t*_8_ = 3.76, Holm-Bonferroni adjusted *p* = 0.03) when participants fixated the slot (0.30 ± .054 s) compared to when they did not (0.378 ± .056 s). No other phase durations were affected by gaze pattern (adjusted *p* > 0.51 in all cases).

**Figure 3.**
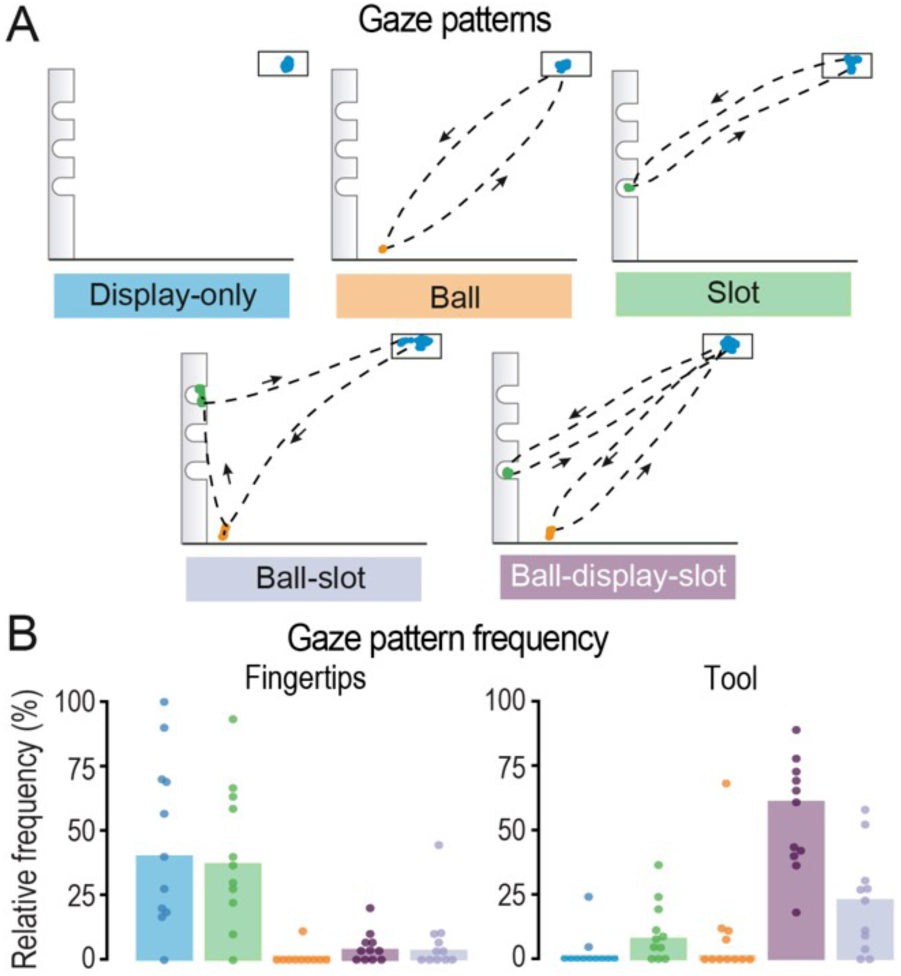
(A) The five distinct gaze patterns observed during dual-task trials are illustrated with data from single trials. (B) Average percentage of each gaze pattern (color-coded as in A) observed across participants in fingertip and tool trials. Individual participants are represented by dots.

In tool trials, participants were much more likely to fixate both the ball and the slot compared to fingertip trials (Fig. 2D). Participants fixated the ball before contact in 88% of trials, and the slot before slot entry in 89% of trials. The probability of fixating the ball peaked towards the end of the reach phase and remained relatively high during the ball approach and most of the grasp phase. Similarly, the probability of fixating the slot peaked towards the end of the transport phase and remained elevated during the slot approach and slot phases. These findings suggest that, in general in tool trials, central vision was crucial for both grasping the ball and inserting it into the slot.

The predominant gaze pattern in tool trials was ‘ball-display-slot’, where gaze shifted from the display to the ball, back to the display, then to the slot, and finally back to the display (Fig. 2D). The second most common pattern was ‘ball-slot’, where gaze shifted from the display to the ball and then directly to the slot before returning to the display (Fig. 3). To examine the relationship between gaze pattern and manual performance, we compared the action phase durations between the ball-display-slot and ball-slot trials. Only participants who exhibited both gaze patterns (*N* = 8) were included in this analysis. We found that the transport phase was shorter (*t*_7_ = 4.71, Holms-Bonferroni adjusted *p* = 0.01) when gaze shifted directly from the ball to the slot (0.32 ± .051 s) compared to when gaze fixated the display between the ball and slot fixations (0.55 ± 0.17 s; *t*_7_ = 4.71). No other phase durations differed significantly (adjusted *p* > 0.22 in all cases). As in the single-task conditions, in the dual task conditions the ball-drop task was performed more slowly with the tool (2.77 ± 0.4 s) than with the fingertips (2.11 ± 0.23 s; *t*_10_ = 5.61, *p* < 0.001; *d* = 1.69).

### Temporal coupling between fixations and contact events

To examine the trial-by-trial coordination between the timing of action fixations and specific kinematic or contact events, we conducted a multiple linear regression analysis to assess how well the onset and offset times of ball and slot fixations could be predicted by the timing of the following six events marking action phases boundaries: (1) start of the reach phase, (2) start of the ball approach phase, (3) time of first ball contact (start of the ball grasp phase), (4) time of ball liftoff (start of the ball transport phase), (5) start of the slot approach phase and (6) time for slot entry (start of the slot phase). Predictors were centered for each participant (by subtracting their mean) to mitigate structural multicollinearity, and participant identity was included as a categorical nuisance factor to account for variance due to differences in task performance speed. Separate regression analyses were conducted for fixation onsets and offsets, as well as for each action landmark (ball and slot) and end-effector (fingertips and the tool).

In both fingertip and tool trials, ball fixation timing was best predicted by ball contact, and slot fixation timing by slot entry. In fingertip trials, ball fixation onset was predicted solely by the time of ball contact (*t*_1,32_ = 2.84; *p* = 0.008), and slot fixation onset was predicted solely by slot entry (*t*_1,158_ = 9.28; *p* < 0.001). Similarly, in tool trials, ball fixation onset was best predicted by first ball contact (*t*_1,243_ = 6.26; *p* < 0.001), and slot fixation onset was solely predicted by slot entry (*t*_1,244_ = 11.2; *p* < 0.001). Comparable results were found for ball and slot fixation offset times in both fingertip and tool trials (p < .004 in all 4 cases). These findings demonstrate a robust temporal coupling between the initiation and termination of action-related fixations (ball and slot) and their associated contact events (first ball contact and slot entry), consistent across both end-effectors.

The timing of action fixation onsets, relative to contact events, showed remarkable consistency across action landmarks and end-effectors. Both ball and slot fixations typically began approximately 0.4 s before ball contact and slot entry, respectively (solid line curves in Figs. 4A). The timing of fixation offsets was also consistent across action landmarks but varied with the end effector used (dashed line curves in Fig. 4A). In fingertip trials, gaze generally shifted away from both the ball and the slot before the contact event, with an average lead time of about 0.15 s. In contrast, in tool trials, gaze typically shifted shortly after the contact event, with an average lag of about 0.05 s.

**Figure 4.**
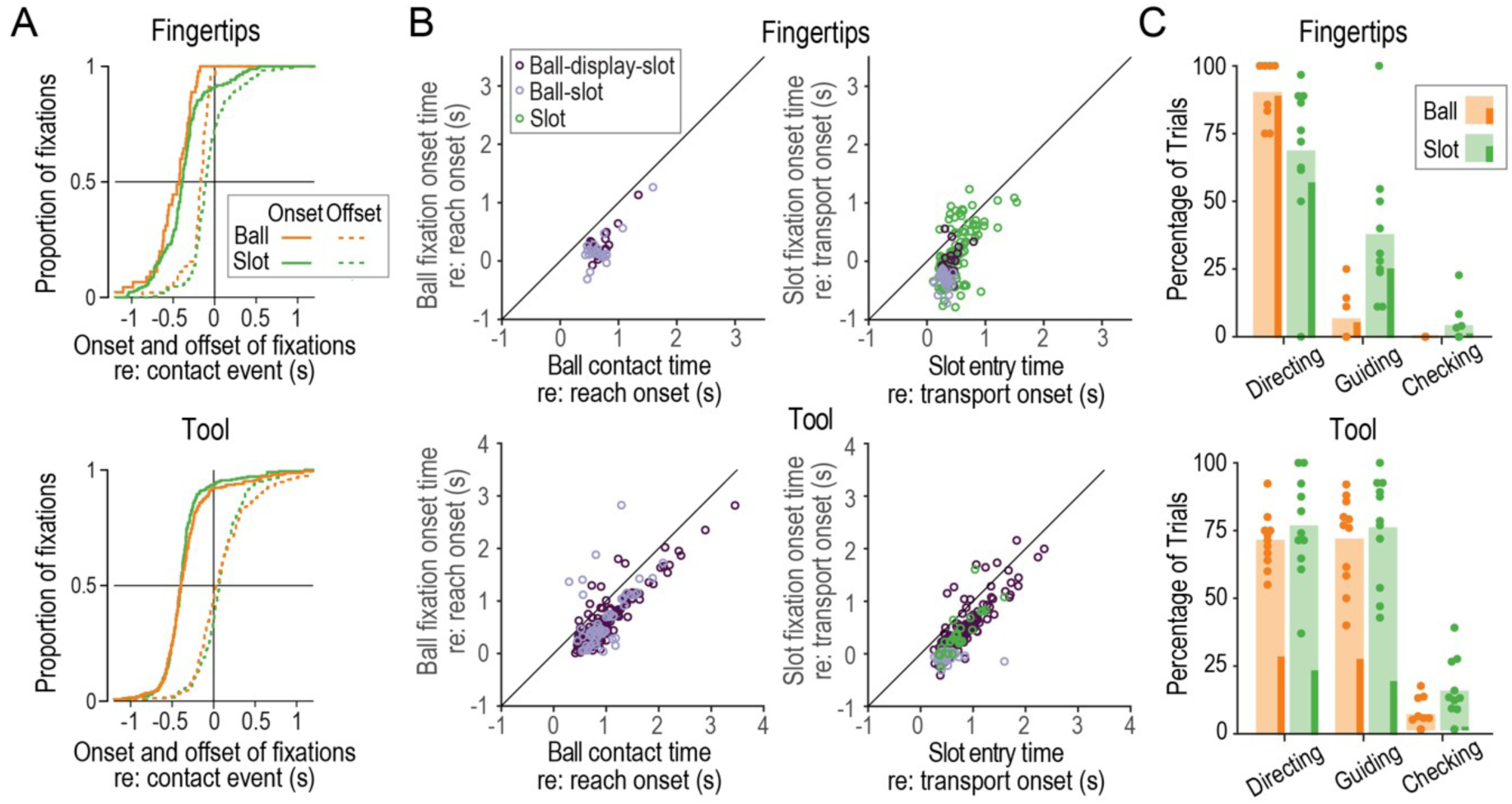
Coordination and classification of action fixations. (A) Cumulative distributions of ball and slot fixation onsets and offsets, aligned to the initial ball contact and slot entry, respectively, in fingertip (top) and tool (bottom) trials. (B) The top row shows, for fingertip and trials, the relationship between ball fixation onset and ball contact times, both relative to reach onset (left), and the relationship between slot fixation onset and slot entry times, both relative to transport onset (right). Dots represent trials from all participants and are colour-coded by gaze pattern. The bottom row shows corresponding data for tool trials. (C) Wide bars show the mean percentage of ball and slot fixations engaged in *directing*, *guiding*, and *checking* across participants in fingertip (top) and tool (bottom) trials. Note that a given fixation could be engaged in more than one function. Circles represent individual participants, except for the circles at zero which represent all participants with zero. The thin bars represent percentages of single-function fixations.

The left column of Fig. 4B shows, for each end-effector, the relationship between the ball fixation onset time and ball contact time, where time is relative reach onset. The right column of Fig. 4B shows the relationship between slot fixation onset time and slot entry time, where time is relative to slot entry. These plots show trials involving the main gaze patterns (i.e., ball-slot, ball-display-slot, and slot trials), which are colour coded. In all four cases (i.e., plots), we observed strong temporal coupling fixation onsets and contact times (r = 0.58 to 0.85, all p values < 0.001), with similar coupling seen across the different gaze patterns. Regression slopes were close to unity (0.89-1.02, mean = 0.98), and intercepts were consistently negative (-0.24 to - 0.47, mean = -0.37). These values confirm that both ball and slot fixations typically began -0.4 seconds before the corresponding contact event, regardless of gaze pattern or end-effector.

### Functional roles of action-related fixations

Although the timing of ball and slot fixations, relative to ball contact and slot entry, was quite consistent, the function(s) served by these fixations (i.e., *directing*, *guiding*, and *checking*) could vary due to intertrial variability in the durations of their corresponding action phases (i.e., reach and ball approach for ball fixations and transport and slot approach for slot fixations). To assess the function, or functions, served by action task fixations, we determined the function of each individual ball and slot fixation. A fixation was classified as *directing* if the ball or slot was fixated for at least 100 ms during the reach or transport phase, respectively. A fixation was classified as *guiding* if the ball or slot was fixated for at least 100 ms between the start of the ball or slot approach phase and the end of the grasp or slot phase, respectively (i.e., the combination of the approach and manipulation phases). Finally a fixation was classified as checking if the ball or slot was fixated for any period of time after the end of the grasp or slot phase, respectively, allowing visual confirmation of the action’s completion (Safstrom et al., 2014). Note that a given fixation could serve multiple functions depending on its timing and duration.

In fingertip trials, ball fixations primarily served a *directing* function, whereas slot fixations often served both a *guiding* and a *directing* function (Fig. 4C, top panel). In contrast, in tool trials, both ball and slot fixations frequently served both *directing* and *guiding* functions (Fig. 4D, bottom panel). Across end-effectors, *checking*.fixations were infrequent. Finally, in fingertip trials, most of the ball and slot fixations served only one function (80% overall), whereas in tool trials, this proportion was lower (48% overall; see thin solid bars within each wide bar in Figs. 5C and D).

**Figure 5.**
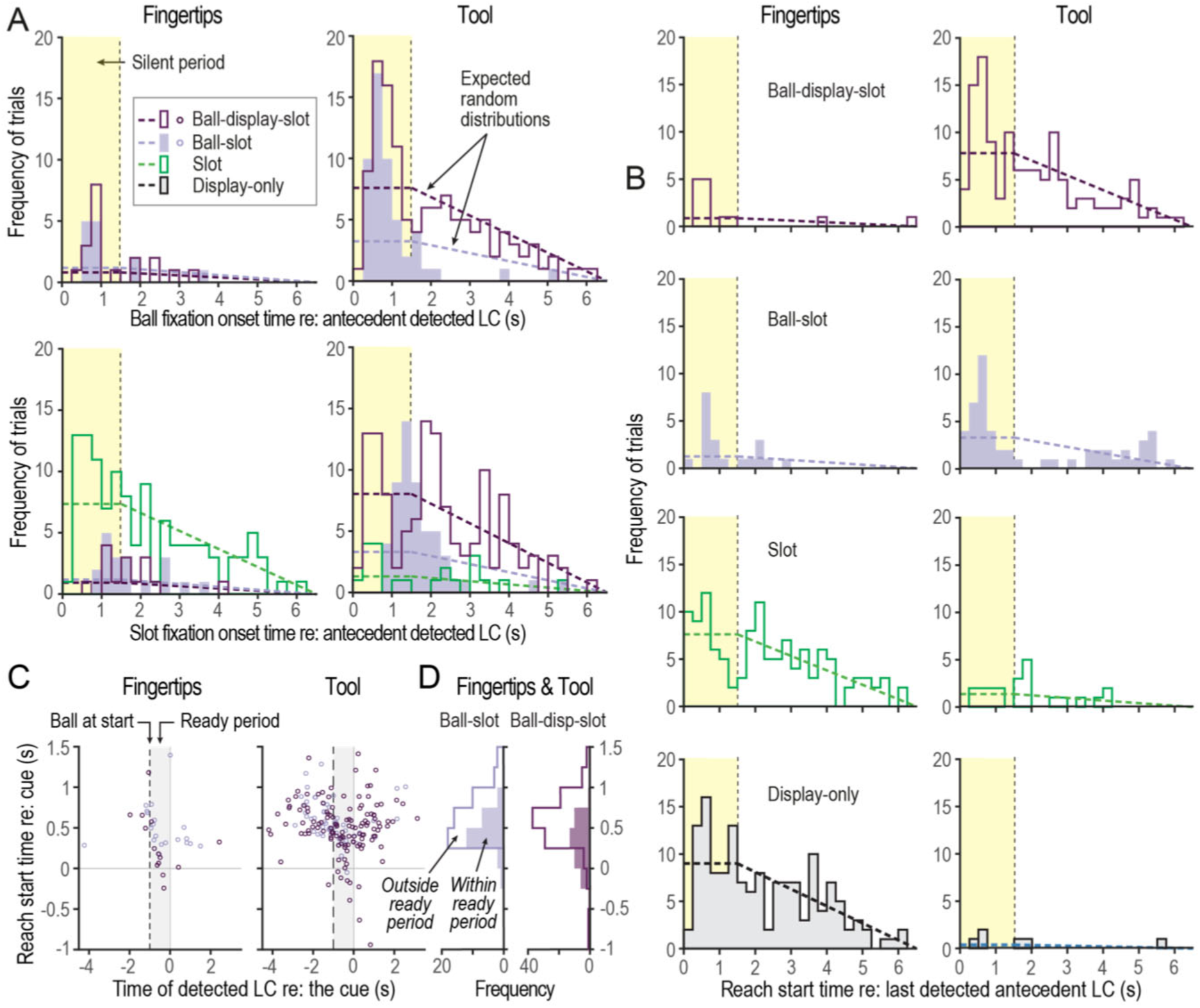
Impact of timing of letter changes (LCs) on gaze fixations and hand movements. (A) The top row shows frequency distributions, combining trials from all participants, of ball fixation onsets-relative to the time of the last detected LC before the fixation onset-in fingertip (left) and tool (right) trials. Yellow regions indicate the silent period. Separate distributions are shown for trials with the main gaze patterns. Dashed lines show expected distributions. The bottom row shows corresponding plots for slot fixation onsets relative to the time of the last detected LC before the fixation onset. (B) Frequency distributions, combining trials from all participants, of reach start times relative to the last detected antecedent LC in fingertip and tool trials. Yellow regions indicate the silent period. Separate plots are shown for the four main gaze patterns. (C) Relationship between reach start time and time of nearest detected LC, both relative to cue onset, in fingertip and tool trials with ball-slot and ball-display-slot gaze patterns. (D) Frequency distributions, combining fingertip and tool trials, of the reach start times in trials with ball-slot and ball-display-slot gaze patterns, aligned with the data shown in (C). Separate distributions are shown for trials in which the LC was within or outside the ‘ready period’ (1 s period prior to the cue).

### Adaptation to temporal structure of monitoring task

We next examined whether participants could learn and exploit the statistical properties of letter changes (LCs) to mitigate competition for gaze between the visual monitoring and manipulation tasks. Our findings reveal that participants adapt their gaze behaviour both directly, by selecting different gaze patterns, and indirectly, by adjusting the timing of their manual actions in response to LC statistics.

In the dual task condition, the interval between LCs was drawn from a uniform distribution spanning 1.5 to 6 s. As a consequence, there was a 1.5 s “silent period” following each LC, during which no new LC could occur. After this silent period, the likelihood of the next LC occurring (i.e., the hazard rate) increased linearly over the subsequent 5 s. On average, participants encountered 1.08 LCs per trial in fingertip trials and 1.41 LC per trial in tool trials. Importantly, LC detection accuracy was high: 88.8 ± 11.8% in fingertip trials and 87.1 ± 9.1% in tool trials (mean ± SD across participants).

#### Gaze allocation relative to letter change timing

To analyze whether LCs influenced gaze allocation to action landmarks, we compared the observed frequency distributions of ball and slot fixations onset times, relative to the most recently detected LC, with the distributions expected under a random model. In this model, fixation onset probability is constant during the silent period (hazard rate = 0) and then decreases linearly over the next 5 seconds as the hazard rate increases from 0 to 1. For each end-effector, we conducted separate analyses for trials exhibiting each of the primary gaze patterns: ball-display-slot, ball-slot, and slot trials.

In both fingertip and tool trials in which participants fixated both the ball and slot (i.e., ball-slot and ball-display-slot trials), the distribution of ball fixations onsets deviated significantly from the expected random distribution (Kolmogorov-Smirnov test, p :S 0.01 in all four cases). Specifically, ball fixation onsets during the silent period occurred more frequently than expected by chance (Fig. 5A, top row).

In tool trials, the choice of gaze pattern was strongly influenced by the timing of the ball fixation relative to the preceding LC. When the onset of the ball fixation occurred during the silent period, participants were equally likely to adopt either the ball-slot or ball-display-slot pattern. However, when the onset of the ball fixation occurred after the silent period, the ball-display-slot pattern was predominantly chosen. That is, participants rarely shifted their gaze directly from the ball to the slot (ball-slot pattern) unless the ball was fixated within the silent period. Conversely, when ball fixation took place after the silent period, participants almost always fixated the display before shifting their gaze to the slot (ball-display-slot pattern).

As expected, in trials with the ball-slot gaze pattern, a peak in the distribution of slot fixation onsets followed the peak in the distribution of ball fixation onsets during the silent period (Fig. 5A, bottom row). This reflects the fact that in 96.4% of these trials, the last LC detected before the ball fixation was also the last LC before the slot fixation. For both fingertip and tool trials, the distribution of slot fixation onsets differed significantly from the expected random distributions (KS test, p < 0.02 in both cases).

Similarly, in trials with the ball-display-slot gaze pattern, a peak in the distribution of slot fixation onsets occurred later than the peak in the distribution of ball fixation onsets, again reflecting trials where the same LC preceded both fixations. In fingertip trials, this slot fixation distribution differed significantly from the expected random distribution (KS test, p < 0.01). In tool trials with the ball-display-slot pattern, an additional earlier peak in the distribution of slot fixation onsets was observed during the silent period, representing trials where a LC was detected while gaze was still on the display prior to shifting to the slot. As a result, the overall distribution of slot fixation onsets did not differ significantly from the expected random distribution (KS test, p = 0.4). In trials where only the slot was fixated (slot pattern), the distribution of slot fixation onsets did not differ significantly from the expected random distribution (KS test, p > 0.06 for both fingertip or tool trials; Fig. 6A, bottom panels).

Together, these results demonstrate that participants adapted their gaze behavior to exploit the statistical properties of LCs, preferentially fixating action targets during low-probability periods to mitigate gaze competition between monitoring and manipulation demands.

#### Timing of manual actions

We found that participants also adapted the timing of their manual actions-specifically, reach initiation-in response to LC statistics. In both fingertip and tool trials in which participants fixated the ball (i.e., the ball-slot and ball-display-slot gaze patterns), the distribution of reach onset times relative to the preceding (or antecedent) detected LC differed from the expected random distribution (K-S test, *p* < 0.05 in all four cases). Reach onsets were biased towards the silent period (top two rows of Fig. 5B), mirroring the bias observed in ball fixation onsets (Fig. 5A). In contrast, in fingertip trials where the ball was not fixated (i.e., the slot and display-only gaze patterns), the distribution of reach onset times did not differ from the random distribution (K-S test, p ≥ 0.4; bottom two rows of Fig. 5B). Because of the limited number tool trials with the slot and display-only gaze patterns, we did not test whether the distributions of reach onsets in these trials differed from the random distribution. These findings suggest that initiating the reach during the silent period was a strategy to reduce the risk of missing LCs.

In the ball drop task, participants typically initiated their reach movement towards the ball in response to the auditory cue signaling the active slot for that trial. These ‘reactive reaches’ generally began about 0.5 s after the cue. However, in a notable proportion of trials, participants initiated their reach in anticipation of the cue, starting either before or shortly after the cue (within 0.5 s). If these ‘anticipatory reaches’ were triggered by a LC occurring shortly before the cue, and accompanied by ball fixation, they could contribute to the greater-than-expected frequency of both reach onsets and ball fixation onsets during the silent period in trials with the ball-slot and ball-display-slot gaze patterns.

To investigate this possibility, we focused on fingertip and tool trials involving ball fixation, and examined the relationship between the timing of reach onset relative to the cue, and the timing of the detected LC relative to the cue. For each trial, we identified the detected LC closest in time to the midpoint of the ‘ready period’, defined as the 1-second interval between the ball returning to its start position and the auditory cue. This LC could occur either before or after the midpoint of the ready period. We observed that most anticipatory reaches-characterized by reach onset times near or preceding the cue-occurred when the detected LC fell within the ready period. This relationship is illustrated in Fig. 5C, which plots reach onset times relative to the cue against the timing of the detected LC relative to the cue. These findings suggest that the decision to initiate an anticipatory reach is closely linked to the detection of a LC during the ready period.

Further support for this idea comes from the frequency distributions of reach onset times relative to the cue, shown in Fig. 5D for trials with the ball-slot and ball-display-slot patterns, Reach onset times occurred earlier when the LC was detected within the ready period compared to when it occurred outside of it. Due to the relatively small number of fingertip trials, fingertip and tool trials were combined for these analyses. Importantly, for both gaze patterns, the distributions within and outside the ready period differed significantly (KS test, *p* < 0.02 in both cases), indicating that the timing of LCs had a distinct influence on reach initiation during the ball drop task.

Overall, these results demonstrate a second strategy by which participants exploited LC statistics to manage gaze competition. That is, in addition to selecting gaze patterns-on a trial by trial basis-to increase the likelihood that gaze could be allocated to the action task with minimal disruption of LC monitoring, participants also adjusted the timing of their reaching movements so that the ball could be preferentially fixated during the silent period.

## Discussion

This study investigated how individuals manage competition for gaze resources when performing an object manipulation task while simultaneously monitoring the environment for visual events. We found that participants employed two complementary strategies to manage this competition. First, they selectively allocated gaze to the action task, prioritizing different gaze functions depending on the haptic capacity of the end-effector. Second, participants exploited temporal regularities of events in the monitoring task, adjusting both gaze allocation and the timing of manual actions to reduce the likelihood that action-related fixations would coincide with visual events.

### Timing, frequency and function of gaze fixations in the ball-drop task

As expected, when the ball-drop task was performed in isolation, gaze was directed to the ball and slot in nearly all trials. Gaze typically arrived at the target (ball or slot) well ahead of the end-effector and departed around the time the end-effected arrived at the target. This behaviour aligns with prior research on eye-hand coordination in visually guided tasks (Land et al., 1999; Johansson et al., 2001; Flanagan and Johansson, 2003; Hayhoe, 2017; Fooken et al., 2021; Tllamperuma and Fooken, 2024).

When participants performed the ball-drop task concurrently with the monitoring task, gaze was intermittently directed to the action task. When the ball-drop task was performed with the fingertips, the ball was fixated infrequently and the slot was fixated in about half of the trials. These fixations were relatively brief (-0.25 s on average) and primarily served a *directing* function, whereby peripheral vision and gaze-related signals support fast, automatic adjustments to the ongoing hand movement (Goodale et al., 1986; Neggers and Bekkering, 2000, 2001; Saunders and Knill, 2003, 2004; Dimitriou et al., 2013; de Brouwer et al., 2018). These results indicate that participants capitalized on the tactile capacity of the fingertips to complete the movement goal (ball grasp or slot entry) without relying on central vision to guide the hand. Tactile information not only guides force adjustments once the ball is grasped, but can also drive kinematic corrections to align the fingertips and ball during grasping or the ball and slot when placing, potentially through distinct neural control mechanisms (Chib et al., 2009). Tmportantly, in manipulation tasks, tactile information enables rapid adjustments (90-120 ms) to both force (Johansson and Flanagan, 2009) and kinematics (Pruszynski et al., 2016; Pruszynski et al., 2018) via fast automatic feedback control processes.

### Tool use and increased visual demands

Using the tool increased the visuomotor demands of the task due to reduced tactile feedback and increased precision demands. We included trial blocks with the tool to alter the balance of gaze demands between the manipulation and monitoring tasks in terms of gaze demands, thereby gaining a broader understanding of how individuals manage gaze competition. Note that tactile sensibility can also be reduced by wearing gloves (Moog et al., 2020) operating in cold conditions (Boada et al., 2016), or peripheral nerve damage (Brink and Mackel, 1987; Jerosch-Herold, 1993). When using the tool, participants almost always fixated both the ball and the slot. These fixations were longer in duration (-0.4 s on average) than those observed in fingertip trials. Moreover, these fixations were typically engaged in both *directing* and *guiding* the end-effector, with the latter involving the use of central vision in deliberate closed-loop feedback control during the final approach to the target (Ballard et al., 1992; Johansson et al., 2001; Land, 2006).

### Temporal anchoring of fixations

Despite differences in functions of action-related fixations between fingertip and tool trials, we found that the onset and offset times of these fixations were most closely correlated with contact events (i.e., ball contact and slot entry for ball and slot fixations, respectively) compared to other events in the ball-drop task such as reach or transport onset. This finding supports the idea that in object manipulation tasks, contact events-which mark sub-goals of the overall task-serve as anchor points for sensorimotor control processes (Johansson and Flanagan, 2009).

### Gaze patterns and task performance

Previous work on eye-hand coordination has often focused on tasks where there is little variation in the sequence of gaze fixation locations. For example, in reaching tasks in which the hand moves to a single target or a series of targets, gaze is almost invariably directed to each target as the hand moves towards it (Epelboim et al., 1995; Wilmut et al., 2006; Bowman et al., 2009; Safstrom et al., 2014). Similarly, when our ball drop task was performed in isolation, participants exhibited a stereotypical gaze pattern. However, when the ball-drop task was performed concurrently with the monitoring task, we found that participants used different gaze patterns across trials. In fingertip trials, two main gaze patterns were observed: “display only” and “slot”, distinguished by whether the participant fixated the slot or kept their gaze on the display throughout the trial. In tool trials, two main patterns were also observed: “ball-slot” and “ball-display-slot”, distinguished by whether gaze was directed to the display between the ball and slot fixations.

Importantly, the choice of gaze patterns was linked to task performance. In fingertip trials, the duration of the slot phase was shorter when participants fixated the slot, and in tool trials, the duration of transport phase was shorter when participants shifted their gaze directly from the ball to the slot, bypassing the display. These results suggest a trade-off between the action and visual monitoring tasks; allocating gaze to the action task improves performance but increases the risk of missing a letter change (LC). However, as discussed below, participants mitigated this trade-off by leveraging the LC statistics when selecting gaze patterns and timing their hand actions.

### Integration of environmental statistics

Our findings support the hypothesis that individuals can exploited the temporal statistics of LCs to guide gaze allocation decisions. In fingertip trials, the decision to fixate the ball was strongly influenced by the LC timing, with almost all ball fixations occurring during the silent period when no LC could occur. Similarly, in tool trials, the decision to fixate the display between fixating the ball and slot was also influenced by LC timing. Specifically, participants often skipped the display when the ball fixation occurred within the silent period, but almost always fixated the display when the ball fixation occurred outside this period. These findings align with previous research demonstrating that human gaze behaviour is sensitive to probabilistic regularities in the environment (Jovancevic-Misic and Hayhoe, 2009). For example, individuals adjust the timing of gaze shifts based on learned temporal statistics of relevant visual events to optimize detection of visual events occurring at different intervals across two spatial locations (Hoppe and Rothkopf, 2016). In visual search tasks, gaze is allocated strategically based on spatial statistics to optimize exploration (Najemnik and Geisler, 2005; Renninger et al., 2007; Eckstein, 2017; Hoppe and Rothkopf, 2019). Our study adds a novel perspective on how environmental statistics influence gaze behaviour by demonstrating that humans can learn and exploit the temporal patterns of externally determined events in the visual environment while concurrently performing an action task that relies on visual guidance.

### Strategic timing of manual actions

Unlike visual monitoring tasks, where event timing is externally determined, in action tasks the timing of ‘events’ that make demands on central vision can generally be controlled or adjusted by the actor. We found support for our hypothesis that individuals can adjust the timing of their manual actions to reduce competition for gaze resources. Specifically, participants modulated reach onset timing so that ball fixations-supporting reaching and grasping-occurred during the silent period more often than expected under a random model. Importantly, this indicates that participants not only learned the statistical properties of LCs but also predicted when and where gaze would be beneficial for action control.

#### Conclusions

This study offers novel insights into the control and coordination of eye and hand movements in real-world scenarios involving competing visual demands. First, we show that when gaze competition arises, participants prioritize key control points-specifically, contact events between the hand, or tool in hand, and target object in the environment-when allocating gaze to the action task. Second, we demonstrate that participants exploit temporal regularities in the external environment to enhance task performance, flexibly adapting both gaze allocation and movement timing to minimize interference between concurrent monitoring and manipulation demands. Together, these findings highlight how humans integrate perceptual and sensorimotor mechanisms to manage visual competition and maintain effective control in complex, dynamic tasks.

## Credit authorship contribution statement

Conceptualization (JF, RSJ, JRF), Methodology (JF, RSJ, JRF), Validation (JF, RSJ, JRF), Formal analysis (JF, RSJ, JRF), Resources (RSJ, JRF), Data Curation (JF, RSJ), Writing - Original Draft (JF, RSJ, JRF), Writing - review & editing (JF, RSJ, JRF), Visualization (JF, RSJ, JRF), Funding acquisition (JF, RSJ, JRF)

## Acknowledgments

This work was supported by a Deutsche Forschungsgemeinschaft (DFG) Research Fellowship to JF (FO 1347/1-1), the Swedish Research Council Project (grant 22209 to RSJ), the Canadian Institutes for Health Research (grant RGPIN/05944-2019 to JRF), and the Natural Science and Engineering Research Council of Canada (grant 156173 to JRF).

